# Smart bionic graspers: background study and design process

**DOI:** 10.1101/2020.09.28.316778

**Authors:** Xinrui Li, Mahmoud Chizari

## Abstract

This paper has focused on reviewing passive bionic grasper and designing a virtual prototype using a computer modelling technique. The main aim of this study is to review existing research and compare their functionalities. This has been followed by introducing a concept design with suitable components. To start the project, generating a clear overview form the most updated and relative knowledge and information on existing designs was the intention of the study. The concept design part of this study uses an iterative process (similar to the Double Diamond Model introduced by Frances et al (2019)) including Discover, Define, Develop and Deliver to complete the design components. Following the concept design process, the detailed theoretical considerations and the features of components selection were then defined. In Develop Phase, the goal was to decide the final design and generate the computer model using SolidWorks. The fourth phase of process was Delivery leading the design evaluation and validation of the generated model or virtual prototype. By completing the process, it is possible to determine the feasibility of the design and the need for improvement. In final stage of the design, a finite element approach using SolidWorks Simulation was performed on the concept. Final design was decided after comparing the concepts in terms of several considerations. A series of simulations were performed on the design to evaluate the durability of the design and extend its functionality. The results showed that the supporting pad was robust enough when dropped down from 1.5 meters height, while the hinge which connecting the finger straps would need further improvement to avoid failure during its practical loading.

## 1 INTRODUCTION

Virtual Reality (VR), featuring with immersion, interaction and imagination, has been developing rapidly in the past 30 years (Wang et al 2019). The popularity of VR gaming has been increased significantly with large amount of investment coming into the field (Chan et al, 2017). Wearable devices, among all the input devices used for Virtual, Augmented, and Mixed Reality (VAMR) and Human-Computer Interface (HCI), have been the most popular choices to realize the interaction of users and computers. Smart and haptic gloves are especially important branch of it as the human hand contributes 90% of the function to the upper limb and is one of the most useful organs (Li et al, 2018). Hence, there is large need in the market for VR-based smart gloves. This led to the settling of the goal of this project, to design a passive but dynamic smart grasper that can be used in gaming industry and other related areas. In following, review on existing product has been performed.

### 1.1 HAND STRUCTURE AND KINEMATIC MODELS

Cerulo et al (2017), Salchow-hommen et al (2019), Dipietro et al (2008), Heo et al (2012), Cobos et al (2008) and Hosseini et al (2018) have pointed out that the bone structure of a human hand includes four fingers consisting of distal, intermediate and proximal phalanges, and 3 joints (Distal Interphalangeal (DIP), Proximal Interphalangeal (PIP) and Metacarpophalangeal (MCP) joints, index finger, middle finger, ring finger and little finger, and thumb consisting of distal and proximal phalanges and 2 joints (Interphalangeal (IP) and MCP). The trapeziometacarpal (TM) joint connected thumb to the carpus and the metacarpocarpel (MCC) joints connected other four fingers to the carpal bone. The CMC joint for thumb enables 2 degrees of freedom (DOF), flexion and extension while the CMC joints for other fingers are only plane joints with 1 DOF, flexion and extension. DIP, PIP and IP can be regarded as hinged joints with 1 DOF, flexion and extension, while MCP joint could be ellipsoidal joint with 2 DOF, *MCP*_*f*_ for flexion and extension, and *MCP*_*a*_ for adduction and abduction, though Cerulo et al (2017) and Salchow-hommen et al (2019) have mentioned TM allows flexion/extension, adduction/abduction and circumduction motion and the wrist allows 2 DOF, ulnar/radial deviation and flexion/extension, 2 DOF, as shown in Figure 1.1(a). The range of motion (ROM) of *MCP*_*f*_ and DIP are between 0 to 90 degrees, the ROM of PIP is between 0 to 110 degrees and the ROM of *MCP*_*a*_ is between −15 to 15 degrees, however, ROM of *MCP*_*a*_ of the middle finger is usually 0 degree (Cerulo et al, 2017). At the resting position of the hand without muscle activation, DIP joints are flexed between 10 and 20 degrees, PIP joints are flexed between 30 and 45 degrees, MCP joints are flexed around 45 degrees and the wrist is extended 20 degrees in neutral radial/ulnar deviation. The movements of joints are limited due to static and dynamic constraints, and therefore, accurate kinematic information from a proper kinematic model is important (Kortier et al, 2014; Kortier et al, 2016). Simplified hand kinematics models are shown in Figure 1.1 (b). Cobos et al (2008) have introduced the main constraints of the finger movements and has showed that there were around 6% error for a 9-DOF simplified hand kinematic model and 13% error for a 6-DOF hand model.

**Figure 1.1:**
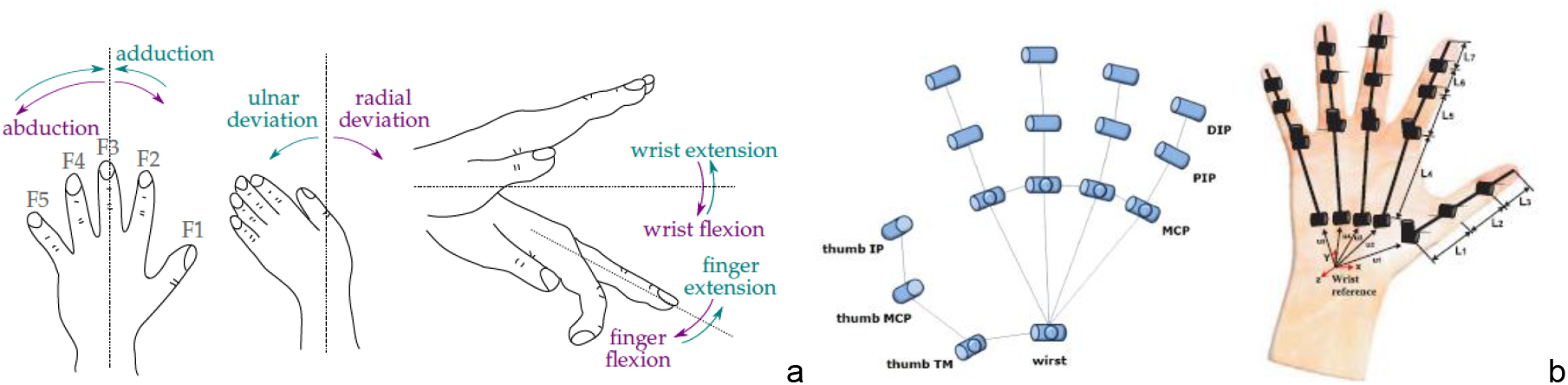
(a) Movements of finger and wrist (Salchow-hommen et al, 2019); (b) Hand kinematic models (Cerulo et al, 2017; Cobos et al, 2008)

### 1.2 EVOLUTION OF SMART GRASPERS AND ITS MAIN CHARACTERISTICS

Since 1940s, researches of haptic display devices have already appeared and in 1965 and the initial concept of VR has been proposed (Wang et al, 2019). There were mainly 3 stages for the development of the human-computer interaction (HCI), desktop, surface and wearable haptics. The interface currently using whole hand to control full virtual hand in the virtual environment has become more convenient, accurate and much easier to perform (Wang et al, 2019). Glove-based systems, as one brunch of wearable devices, initiated in 1970s, and then merged with the robotic, haptic and VR technologies for a new era in recent years (Dipietro et al, 2008). This has significantly expanded the scope of applications for the wearable robotic devices, especially the robotic gloves. Data Gloves were the first generation of glove-based systems since 1970s, although they tended to hold constraints originated from glove cloth and were lack of customization and complicated calibration process (Dipietro et al, 2008; Kortier et al, 2014). Such disadvantages then encouraged further developments, according to Shahid et al (2018). Haptic glove, as one of the most typical wearable haptics, is able to allow users to interact with the computer in variable easy and direct ways (Wang et al, 2019). The latest robotic designs showed that soft materials are much more preferred nowadays than hard materials (Shahid et al, 2018).

Haptic display of HCI has been explained by Wang et al (2019) as an interface for realizing signal communications and interaction between human and computer. It enables users to control virtual objects on the display screen and receive real-time haptic feedback. As Figure 1.2 shows, any movement made by the user to interact with a virtual object through an interface device will be detected and transmitted to the virtual environment. Haptic feedback from the virtual environment will be translated back to the device and then sensed by the user. Compared to the haptic sensation obtained from various ways in real, physical life (such as temperature, humidity and roughness), most of current HCI systems can only provide visual, auditory and simple haptic feedback (i.e. forces or vibrations) (Wang et al, 2019).

**Figure 1.2:**
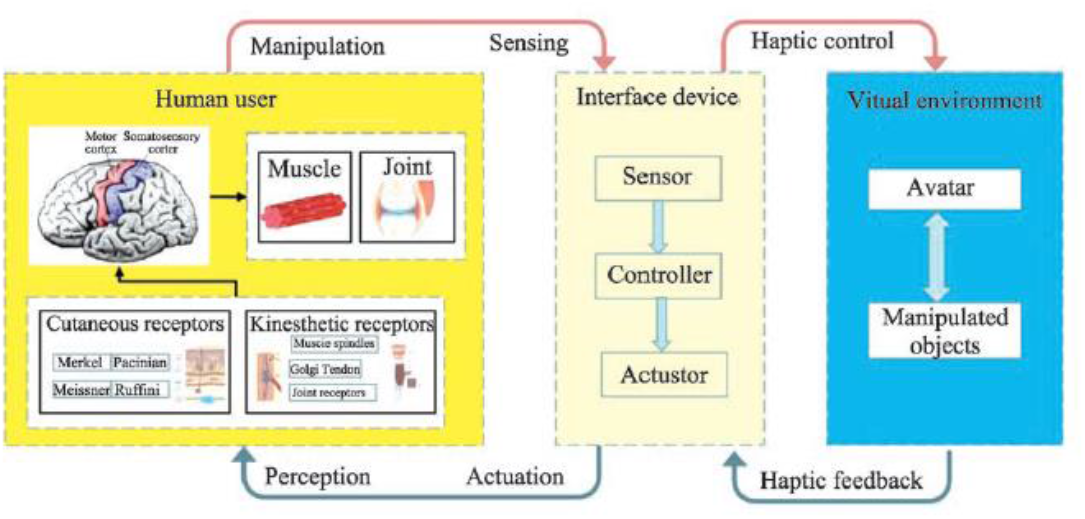
Haptic rendering (Wang et al, 2019)

Zheng et al (2018) has summarized that robotic gloves with force-feedback can be divided into two categories by actuation solutions, active and passive robotic gloves. The active ones, which can provide active control/feedback to the hand, usually employ electric, hydraulic or pneumatics actuators. Safety is a general concern about active force feedback systems, especially when the system is out of control (Hirata et al, 2011, Zheng et al, 2018). Passive actuation solution, on the other hand, would not face such concerns, although the disadvantages of it would be that no actuation can be generated if the hand stays motionless.

One of the most challenging problems for designing is to track the hand motion accurately in real time. The recognizing accuracy of current data glove systems seems to be enough, however, for real-time online gaming, precision and high recording and transmitting rate should be the priority (Salchow-hommen et al, 2019). Table 1.1 listed 5 main types of tracking solution systems. In terms of the sensor placement, Dipietro et al (2008) have demonstrated that sensors can either be sewn in the clothes or be mounted on the structure of gloves without clothes. 2 typical sensor distributions are shown in Figure 1.3, in which 3 sensors were placed on each finger (2 on the thumb) in (a), while flex sensors were used in (b). Similar applications can be also found in the studies by Weber et al (2016), Shin et al (2016) and Bhardawaj et al (2018).

**Table 1.1:** 5 main hand motion tracking systems (Salchow-hommen et al, 2019)

| Type | Advantages | Drawbacks |
| --- | --- | --- |
| Optical tracking with markers | High accuracy | Expensive; Complicated setup |
| Optical tracking without markers | Provide high DOFs | Not fully reliable; Not working in case of occlusions |
| Flex-sensors/bend-sensors tracking | Good measurements; Easy setup | Continuous bending reduces lifetime; Easier to break |
| IMU tracking | Quick setup | Not working well in full-of-vibration systems |
| Rotational-sensors tracking | Reliable | Cost and space consuming |

**Figure 1.3:**
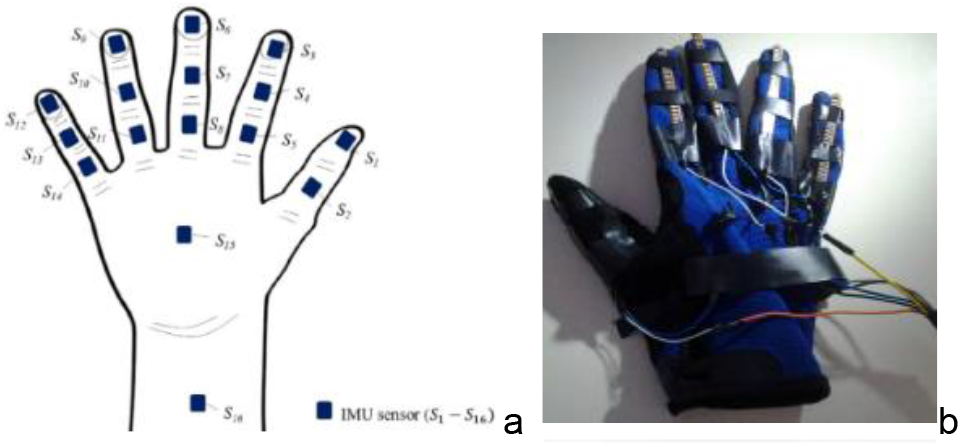
Sensor distribution (a) (Lin et al, 2019); (b) Flex sensors placed on each finger (Silva et al, 2013)

Several processing/controlling systems have been proposed by Das et al (2016), Wilk et al (2018), Zheng et al (2018), Frawan et al (2018) and Zou et al (2020), etc. The workflows of these systems were similar: a micro control unit would collect raw data from all of the sensors placed on the smart grasper and transmit the readings in real-time, usually via Bluetooth, to the sink (usually the computer) for further segmentation, recognition and assessment.

One challenge is the complexity of the body shapes and functions, such as the dexterousness of the hand, thus, existing smart glove designs have followed the ‘trade-off’ rule in order to obtain the best balance of each feature (Ma and Ben-Tzvi, 2015). It was studied by Lariviere et al (2010) that wearing gloves may lead to increase of up to 21% of the forearm muscle activation due to stiffness, which could affect the wearing experience. Hence, the comfort of a smart grasper would be important. Besides, Zheng et al (2011) stated that 2500 to 3000 grasping motions were usually performed during an 8h working period in a day, indicating durability should be also one of the key considerations. User-friendly and real-time have become the leading needs currently, which can be translated as ergonomic, lightweight, effective and comfortable. 10 glove-based systems have been compared in Table 1.2, which showed the most typical technologies available for the glove-based systems, and would help with drafting the design.

**Table 1.2:**
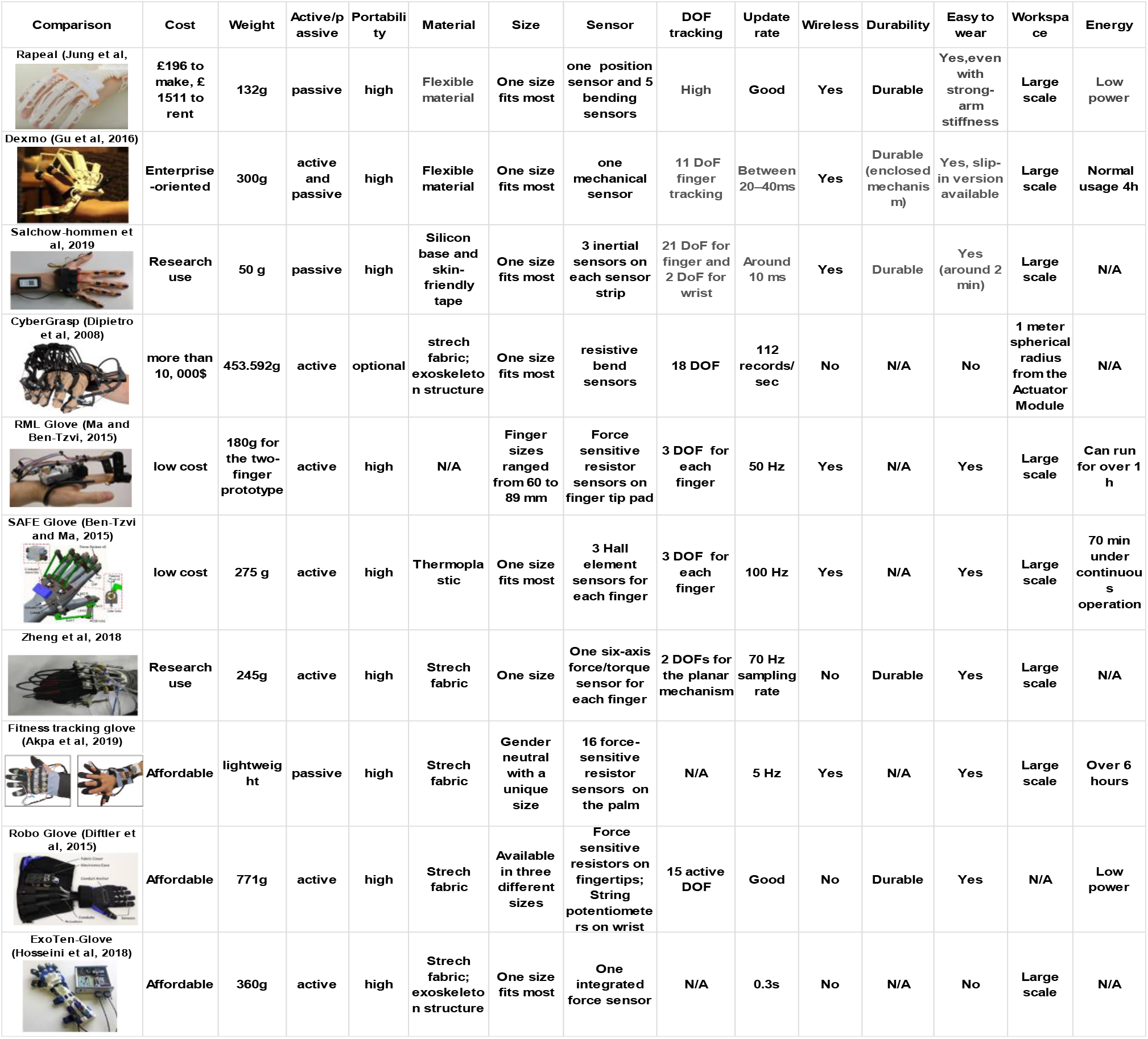
Comparison of 10 typical glove-based systems in recent years for research/commercial use

| Comparison | Cost | Weight | Active/passive | Portability | Material | Size | Sensor | DOF tracking | Update rate | Wireless | Durability | Easy to wear | Workspace | Energy |
| --- | --- | --- | --- | --- | --- | --- | --- | --- | --- | --- | --- | --- | --- | --- |
| Rapeal (Jung et al, | £196 to make, £1511 to rent | 132g | passive | high | Flexible material | One size fits most | one position sensor and 5 bending sensors | High | Good | Yes | Durable | Yes, even with strong-arm stiffness | Large scale | Low power |
| Dexmo (Gu et al, 2016) | Enterprise-oriented | 300g | active and passive | high | Flexible material | One size fits most | one mechanical sensor | 11 DoF finger tracking | Between 20–40ms | Yes | Durable (enclosed mechanism) | Yes, slip-in version available | Large scale | Normal usage 4h |
| Salchow-hommen et al, 2019 | Research use | 50 g | passive | high | Silicon base and skin-friendly tape | One size fits most | 3 inertial sensors on each sensor strip | 21 DoF for finger and 2 DoF for wrist | Around 10 ms | Yes | Durable | Yes (around 2 min) | Large scale | N/A |
| CyberGrasp (Dipietro et al, 2008) | more than 10,000\$ | 453.592g | active | optional | stretch fabric; exoskeleton structure | One size fits most | resistive bend sensors | 18 DOF | 112 records/sec | No | N/A | No | 1 meter spherical radius from the Actuator Module | N/A |
| RML Glove (Ma and Ben-Tzvi, 2015) | low cost | 180g for the two-finger prototype | active | high | N/A | Finger sizes ranged from 60 to 89 mm | Force sensitive resistor sensors on finger tip pad | 3 DOF for each finger | 50 Hz | Yes | N/A | Yes | Large scale | Can run for over 1 h |
| SAFE Glove (Ben-Tzvi and Ma, 2015) | low cost | 275 g | active | high | Thermoplastic | One size fits most | 3 Hall element sensors for each finger | 3 DOF for each finger | 100 Hz | Yes | N/A | Yes | Large scale | 70 min under continuous operation |
| Zheng et al, 2018 | Research use | 245g | active | high | Stretch fabric | One size | One six-axis force/torque sensor for each finger | 2 DOFs for the planar mechanism | 70 Hz sampling rate | No | Durable | Yes | Large scale | N/A |
| Fitness tracking glove (Akpa et al, 2019) | Affordable | lightweight | passive | high | Stretch fabric | Gender neutral with a unique size | 16 force-sensitive resistor sensors on the palm | N/A | 5 Hz | Yes | N/A | Yes | Large scale | Over 6 hours |
| Robo Glove (Diftler et al, 2015) | Affordable | 771g | active | high | Stretch fabric | Available in three different sizes | Force sensitive resistors on fingertips; String potentiometers on wrist | 15 active DOF | Good | No | Durable | Yes | N/A | Low power |
| ExoTen-Glove (Hosseini et al, 2018) | Affordable | 360g | active | high | Stretch fabric; exoskeleton structure | One size fits most | One integrated force sensor | N/A | 0.3s | No | N/A | No | Large scale | N/A |

**Table 1.3:**
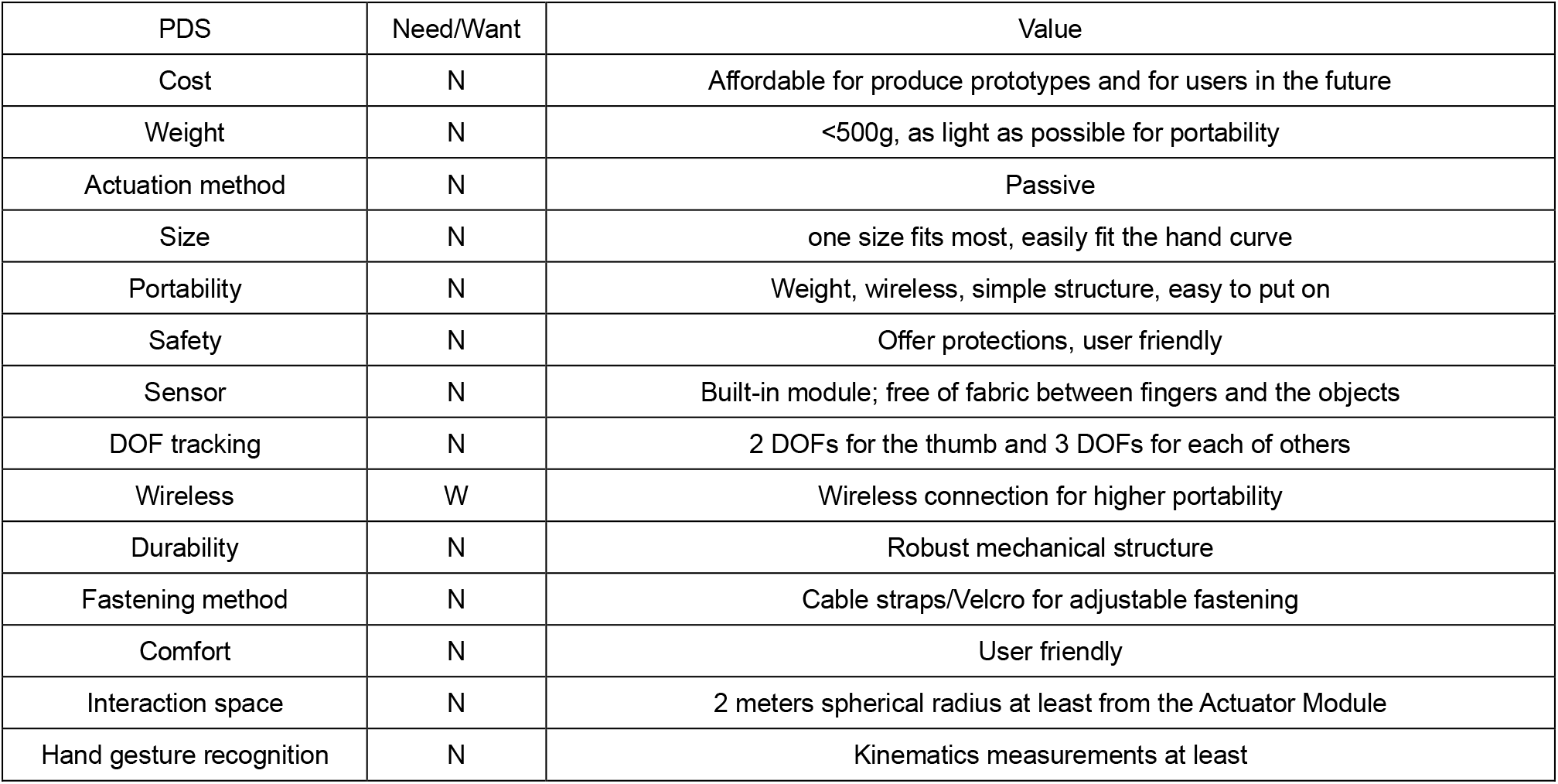
Product design specification (PDS)

| PDS | Need/Want | Value |
| --- | --- | --- |
| Cost | N | Affordable for produce prototypes and for users in the future |
| Weight | N | <500g, as light as possible for portability |
| Actuation method | N | Passive |
| Size | N | one size fits most, easily fit the hand curve |
| Portability | N | Weight, wireless, simple structure, easy to put on |
| Safety | N | Offer protections, user friendly |
| Sensor | N | Built-in module; free of fabric between fingers and the objects |
| DOF tracking | N | 2 DOFs for the thumb and 3 DOFs for each of others |
| Wireless | W | Wireless connection for higher portability |
| Durability | N | Robust mechanical structure |
| Fastening method | N | Cable straps/Velcro for adjustable fastening |
| Comfort | N | User friendly |
| Interaction space | N | 2 meters spherical radius at least from the Actuator Module |
| Hand gesture recognition | N | Kinematics measurements at least |

### 1.3 PRODUCT DESIGN SPECIFICATION

As mentioned above, a satisfied design is based on the trade-off of different parameters. A high-performance smart glove would require large number of different mechanical, electrical and functional characteristics. PDS below was completed based on the literature survey.

## 2 BASIC CONCEPTS SETUP

Before building up the CAD model of the smart grasper, it is necessary to draw a CAD model of hand first, of which the dimensions can be the references for the smart grasper model later. The dimensions of the left-hand model were decided according to (Yu et al, 2013). The simplified model shown in Figure 2.1 mainly imitated the main joints between phalanges, fingers and palm, and palm and wrist, which enabled the main movements by the joints, including finger flexion and finger extension and wrist rotation.

**Figure 2.1:**
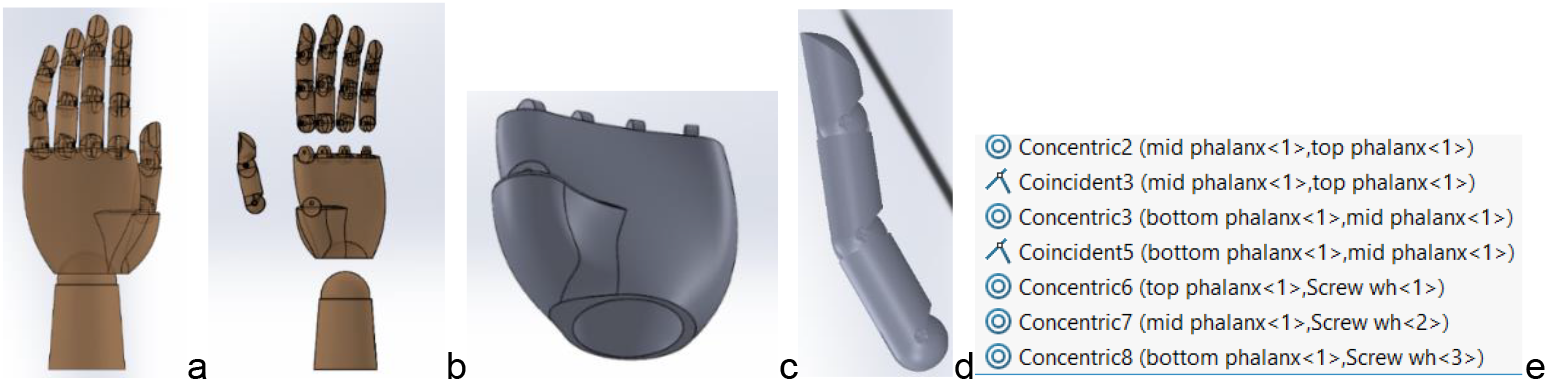
(a) Front view of the hand model; (b) Exploded view of the hand model: 5 fingers, palm and wrist pedestals; (c) The palm pedestal; (d) Side view of the middle finger; (e) Mates of 3 phalanges for the middle finger model

According to Table 1.2, soft clothes have been used frequently in the design of robotic glove. Hence, Concept 1 was designed using soft material. As in Figure 2.2 (a), it consisted of a padding area covering the wrist and the back of the hand with soft fabric for printed circuit board (PCB), assembled with the battery charging module and all the electronics and attached to the finger mechanisms. Velcro band was used on the wrist for fastening. The soft fabric could perfectly match the curve of the hand during motions. Each finger mechanism included one flexible tendon-like strip with 3 Hall effect sensors (2 on the thumb) for measuring the joint angles, and a fingertip cap connected to the strip. The finger flexion and extension, abduction and adduction were enabled due to the flexibility of the strips. For an additional force feedback mechanism, passive actuation design can be assembled into the PCB, connected to 5 tendon cables using tendon-like Dyneema cables (Jeong and Cho, 2016), with the advantages of high-flexibility, high-strength and lightweight material, so that it can transmit necessary passive force along the finger. The material and components selection here meant low cost and lightweight. Besides, Velcro bands allowed the users to easily put on the smart grasper and adjust the tightness. However, the drawbacks are that fingertips, as the most critical area for the tactile sensing function, were covered by fabrics, and with the fabric being in between hand and objects, uncomfortable friction might appear. Besides, disinfection of the grasper might be a problem.

**Figure 2.2:**
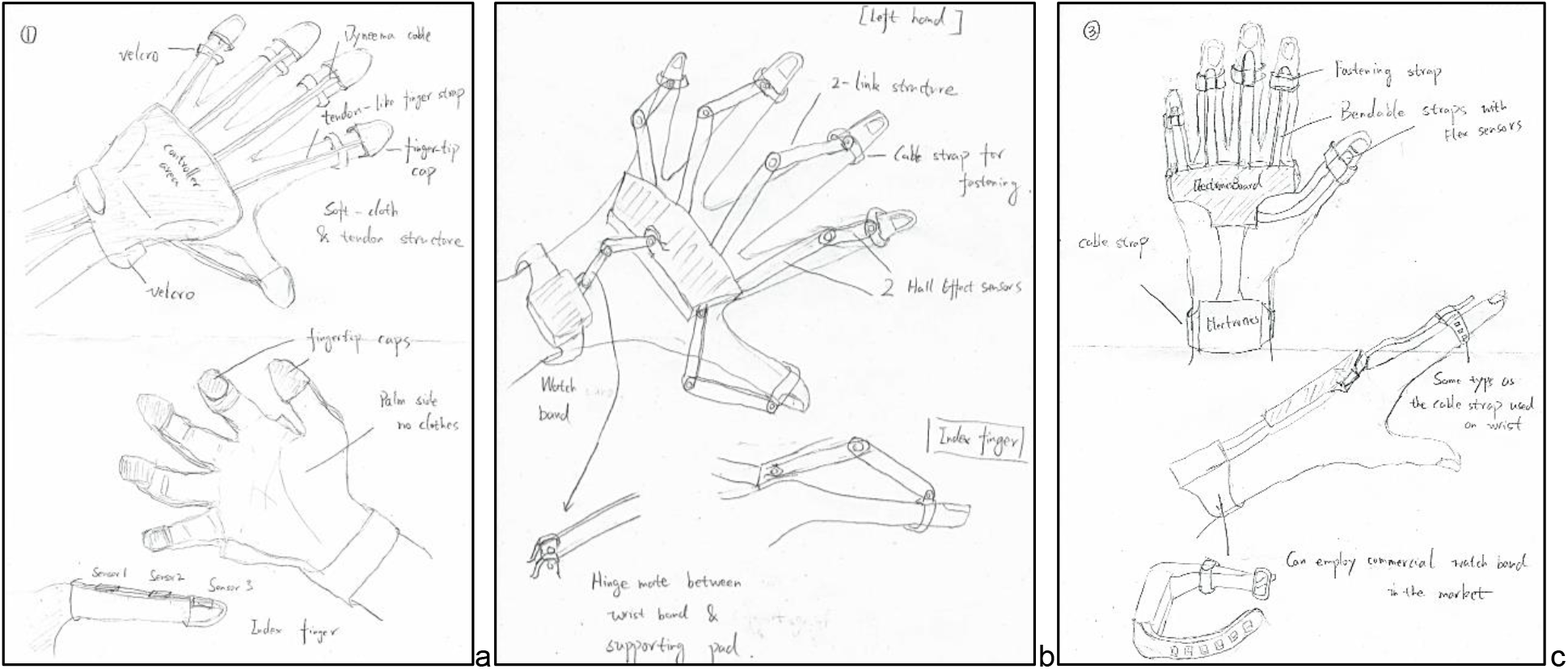
Sketches of (a) Concept 1; (b) Concept design 2; (c) Concept design 3

Concept 2, as in Figure 2.2 (b), has employed no soft clothes. 2 Hall-effect sensors were used on the 2-link cables to be fastened with the fingers. The wrist band structure was same as normal sports bands. The supporting pad can be customized to match the specific curve of the hand for higher comport level, with Thermoplastic material due to its advantages of being anti-sweat and lightweight. It also provided protection to the electronics in it. Flexible ring-shape straps on the finger strap using polyurethane rubber was used for fastening each finger due to its flexibility, water-resistance ability, low cost and high durability. An IMU sensor could be placed in wrist band for monitoring the forearm and wrist movements and Arduino Uno board would be inside of the supporting pad for processing. Passive force-feedback mechanism can be added along the 2-link finger cable using Dyneema cables (Jeong and Cho, 2016) so that the abduction/adduction movements of finger would not be affected, but the extension/flexion movements could be controlled if the mechanism was triggered. This concept has minimized the material/obstruction between hand and objects. The components selections were all lightweight and easy to be cleaned by simply wiping over and can fasten with different hand sizes. 2 sensors for each finger were cheaper than 3 for each finger in Concept 1, but this might lower the accuracy of kinematic measurements. The abduction/adduction movements may be limited by the finger cable. The pad customization may increase the total cost as well.

Concept 3 (Figure 2.2 (c)), similar to Concept 2, has employed an exoskeleton structure without any soft clothes. This was inspired by the design of Rapael Glove (Jung et al, 2017). Same supporting pad and wrist band designs as in Concept 2 was used. It is also optional for the supporting pad to be customized or to use a general curvy structure that fits most of the hand sizes to lower the cost. Same cable straps for fastening were used on both wrist and fingers so that the tightness can be adjusted. The main difference was that one flex sensor were placed on each elastic finger strap, which was attached to the supporting pad, for kinematics measurements. The elastic finger straps would not affect finger movements as the bending range of the flex sensor was up to 180 degrees pinch bend. The total cost of sensors might be slightly higher. Besides, pursuing additional feedback control may increase the complexity.

The comparison of 3 concepts was illustrated in Table 2.1 in terms of 6 objectives. The weight of each concept for each objective were comparably decided based on the relative contents from the literature survey.

**Table 2.1:** Comparison of 3 Concept designs

| Objective | Weight | Concept 1 |  | Concept 2 |  | Concept 3 |  |
| --- | --- | --- | --- | --- | --- | --- | --- |
| Comfort | 0.2 | Lightweight; friction with clothes;<br>covered fingertip | 1 | lightweight, fingertip free<br>of covering | 3 | Lightweight; fingertip<br>free of covering | 3 |
| Hygiene | 0.1 | Soft clothes, | 1 | easily wipe | 4 | easily wipe | 4 |
| Adjustable | 0.1 | not for fingers | 1 | yes | 4 | yes | 4 |
| Accuracy | 0.2 | 3 sensors for each finger | 4 | 2 sensors | 2 | One flex sensor | 4 |
| Complexity | 0.2 | low | 4 | high | 2 | relatively low | 3 |
| Cost | 0.2 | low | 4 | Relatively low | 2 | Relatively low | 3 |
| SUM | 1 | 2.8 |  | 2.6 |  | 3.4 |  |

## 3 COMPUTER MODEL GENERATION AND SIMULATIONS

### 3.1 CAD MODEL

With the highest result of 3.4, Concept 3 has surpassed the other two to be the best choice. The computer models generated for the final design were presented in Figure 3.1–3.3. One end of the supporting pad was designed to connect the wrist band with elastic, soft-material band in order to provide enough length/space for and to avoid restrictions on the wrist movement. The shape of the wrist band has generated from the most common shape of commercial sports bands.

**Figure 3.1:**
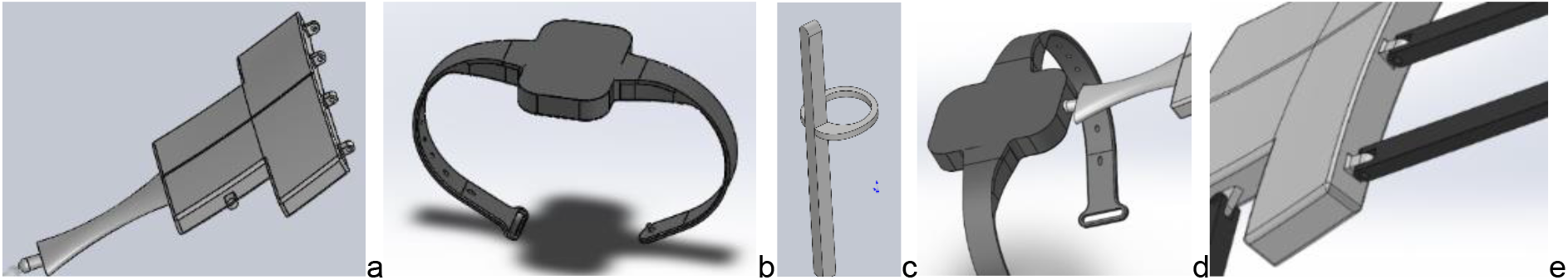
Components design: (a)supporting pad; (b)wrist band; (c)finger strap; (d)Concentric mate between wrist band and supporting pad;(e) Hinge mate between finger strap and supporting pad

**Figure 3.2:**
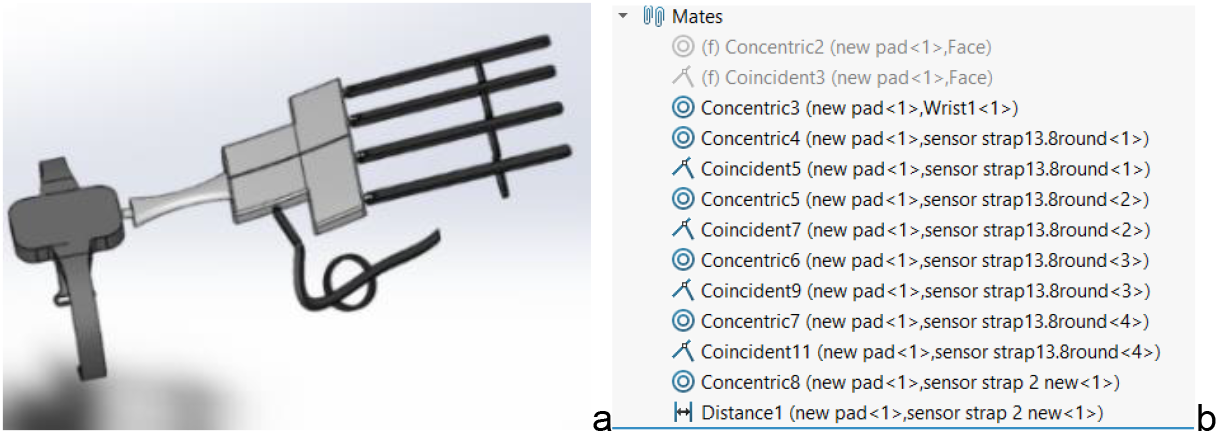
Assembly of the smart grasper model (a) and the mates between each component (b)

**Figure 3.3:**
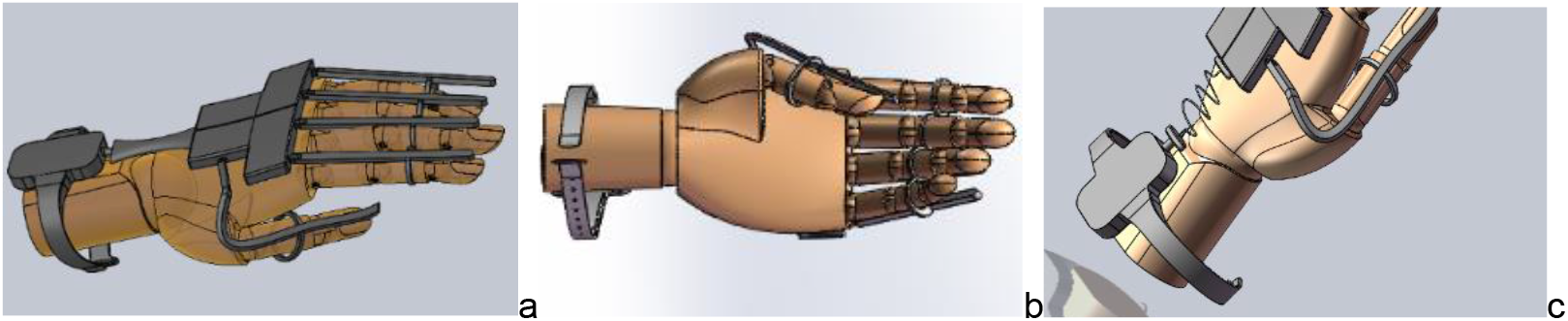
Front (a) and rear (b) view of assembled glove on a hand model; (c)modified smart grasper CAD model

Unlike in real world, it was impossible for SolidWorks to simulate the bending property of a material, such as a thin piece of copper foil or a flexible polyurethane rubber strap. Moreover, this smart grasper was designed as one-piece prototype, however, most of the movements, especially those done by fingers and the wrist, would not be able to be realised if the CAD model is generated as one-piece, thus in the CAD model, the grasper was separated into 7 components, wrist band, supporting pad, and five finger straps. In order to illustrate the ideas of design, mates between different components became very critical. The concentric mate between wrist band and supporting pad was illustrated in Figure 3.1(a) as the material chose in the final design was stretchable, elastic bands, which cannot be directly presented in the CAD model without restricting the wrist movements. Hinge mates were applied between finger straps and supporting pad to enable the flexion and extension of fingers in Figure 3.1(e). However, this may obstruct the abduction and adduction of fingers. Considering the initial prototype of this smart grasper design would mainly focus on the grasp motion, this was acceptable for current stage. Adjustable finger straps have been simplified into closed, round and thin straps enclosing the second phalange of each finger to reduce the workload during limited time without influencing final assembly, demonstration and simulations. The wrist band in Figure 3.3 (a) has not enclosed around the wrist tightly as it was for illustrating the shape. For a better illustration of the availability of wrist extension and flexion on the smart grasper CAD model, it was then decided to change the connection structure between one end of the supporting pad and the wrist band in former drawings into a zig-zag structure shape using the swept-boss/bass feature in SolidWorks in Figure 3.3 (c) to show how the smart grasper was designed to provide enough length/space for and to avoid restrictions on the wrist movement.

### 3.2 DYNAMIC DROP TESTS

It was mentioned by Bocker and Malmstrom (2018) that no standardised test methods were available for soft exoskeleton-structure graspers yet. Thus, tests generated to evaluate the performance should depend on the demands and functionalities of the design itself. Considering only virtual evaluations can be performed in the current situation, this section has been divided into 2 parts, dynamic drop tests for durability validation of the supporting pad and static tests conducted on the index finger using SolidWorks Simulation. The CAD model used here was the first computer generation model in the last section because for both models, only the illustration methods of the wrist-pad connection band were different, which was not the focus of the simulations, and the delicate structure of the swept connection in the second model may cause meshing problems. All the screenshots of the results here were deformed results.

From Chapter 3.1, it can be seen that the match of smart grasper to the hand was satisfied. This means the requirement on comfort might be met. It was mentioned by Gu et al (2016) that a robust product should withstand the hit from other impacts or being dropped down to the ground. Therefore, in this part, the durability of the design has been tested using a dynamic FEA study in SolidWorks for evaluating the effect of the impact of a part or an assembly with a rigid or flexible planar surface. All Components except supporting pad employed elastic soft materials, polyurethane rubber. The supporting pad, as the part connecting to the finger straps and wrist band, and providing protection to the electronics built in it, would be the most critical component for drop test. For time efficiency concern, only the supporting pad has been studied. The first step is to apply the material, Acrylonitrile Butadiene Styrene (ABS), which is the most commonly used thermoplastic material. Next, connections were defined. Considering hinge mates were mainly applied in this assembly, Global Contact would fit here. Standard mesh with coarse quality was unable to be generated, probably because the structure in some area was too subtle for the selected mesh size. In Figure 3.4(a), mesh density has been increased slightly with the standard mesh parameter and successfully generated, following with two more mesh generation with higher density and with curvature-based mesh. The last mesh generation gave the finest results and therefore was used for the simulation as curvature-based mesh solution benefits more from the multi-core processors for using smaller elements at the curvature are, while standard mesh has the advantages on time efficiency. Figure 3.4(d) gave the drop direction from1.5 meters high (from the lowest point) with reference plane. The reason of choosing 1.5m height for the drop test was that this smart grasper was designed for VR gaming, which would be used by users when sitting or standing and the working height range of the grasper would be within the range of 0m to around the height of a normal person.

**Figure 3.4:**
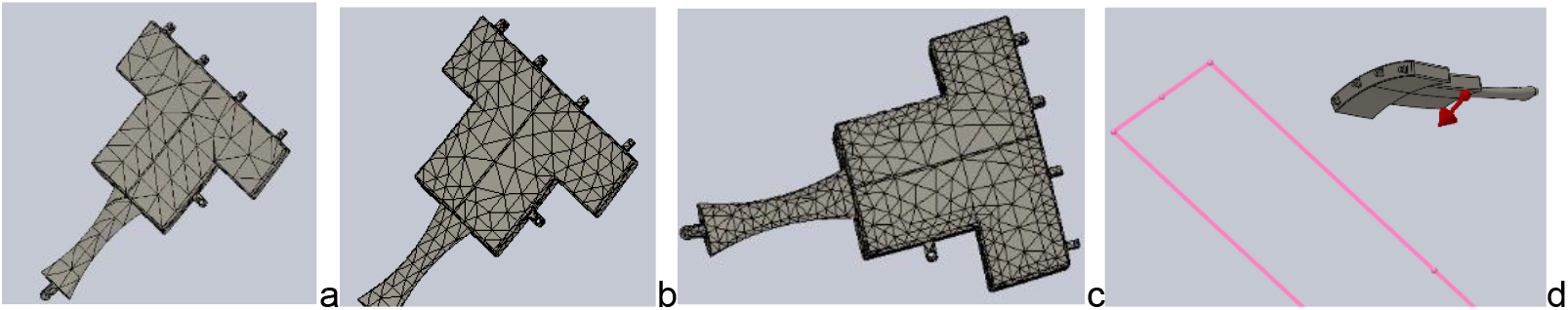
Mesh overview: initial coarse mesh (a); medium size mesh (b); optimised mesh (c); Defining boundary condition for drop test (d)

After all the properties determined, the study was run. One notification was popped up claiming that the acute-angled elements in the CAD model may lead to failure of the simulation. The test has been continued, but it is necessary to note the accuracy of the simulation might be influenced. This analysis has lasted over 7h and therefore, for the consideration of time efficiency and possible crush down of the software during long period working, no more model with even finer mesh has been tested. Final results were shown in Figure 3.5. The Von Mises stress here was lower than the yield strength of the material, 4.8×10^7 N/m^2, thus, yielding would not occur. The strain value was relatively high on the curve, small angle and edge area, which means these areas would be more likely to deform.

**Figure 3.5:**
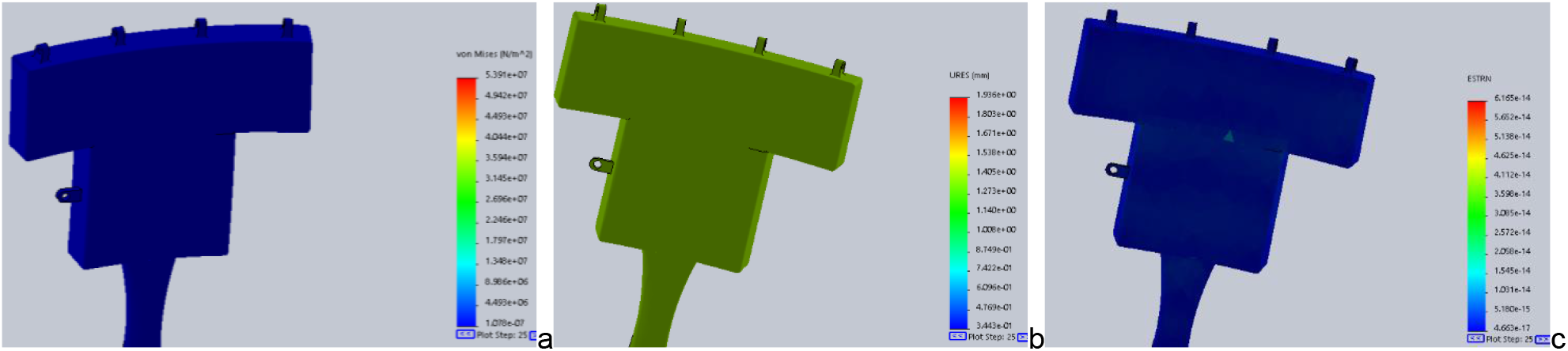
Simulation result: von Mises stress (a); displacement (b); strain (c)

One more drop test has been performed by changing the reference plane for the gravity. It was changed from the flat face of supporting pad dropped to the ground first to the side edge dropped to the ground first. Results were shown in Figure 3.6. The Von Mises Stress indicated no yielding. The displacement value has increased to around 1.4 mm. The strain value at the middle edge tended to be larger than other area. Its durability under this condition has also been proved to be satisfied.

**Figure 3.6:**
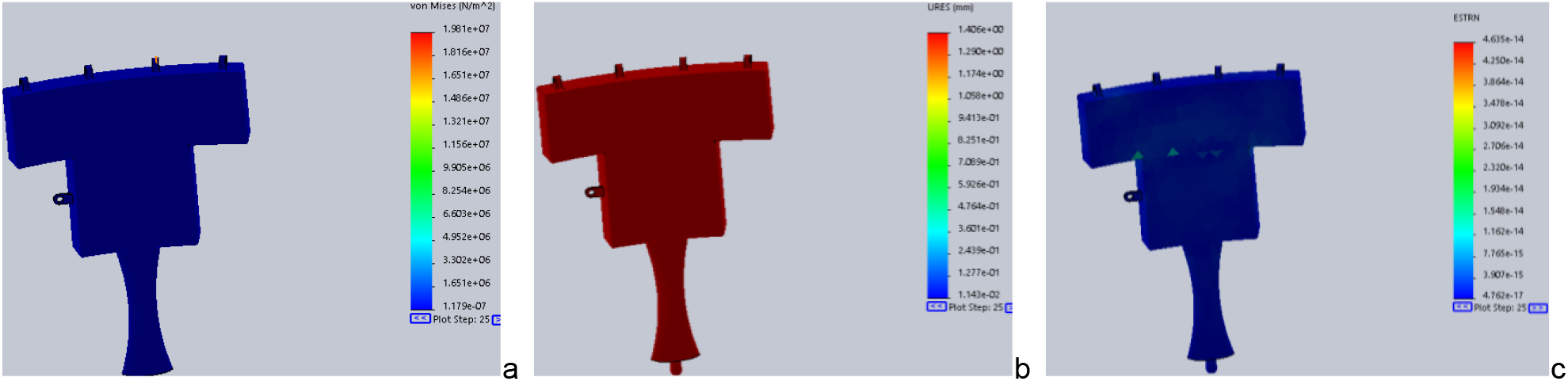
Simulation result after change the gravity’s reference plane: Von Mises stress (a); displacement (b); strain (c)

### 3.3 STATIC SIMULATION

Static evaluation of the model using SolidWorks Simulation was completed for evaluating one of the most frequent tasks for hand, holding the objects (Zheng et al, 2011). In et al (2011) studied that average force required for performing daily activities was approximately 18N. For time efficiency concern, the simulations were mainly run on the Polyurethane rubber index finger strap with yielding strength of 7.89×10^6 N/m^2 and ABS plastic supporting pad. Connections were defined as global contact as only hinge mate was applied. Considering the supporting pad would stay on the back of the hand stably while the smart grasper was in task, the bottom faces of the supporting pad have been chosen for the fixtures, and external loads, 18N, was applied to the bottom face of the index finger strap, as in Figure 3.7(a). Mesh was set up to be the finest. The finger strap was hinged with the supporting pad. Large displacement theory was applied by the solver for more accurate results and avoid running failure. The results were shown in Figure 3.8. Yielding appeared at the area with the hinge mate. More than 35mm displacement happened at the fingertip area while the value gradually decreased from the tip to the end along the strap. However, the results may not mean the failure of the design. As mentioned above, structure with such mate methods was created to illustrate the idea of the final design. The results might be differed if this can be expressed in a better way. Unfortunately, due to the time limits, no more simulation or better presentation of the design can be illustrated here.

**Figure 3.7:**
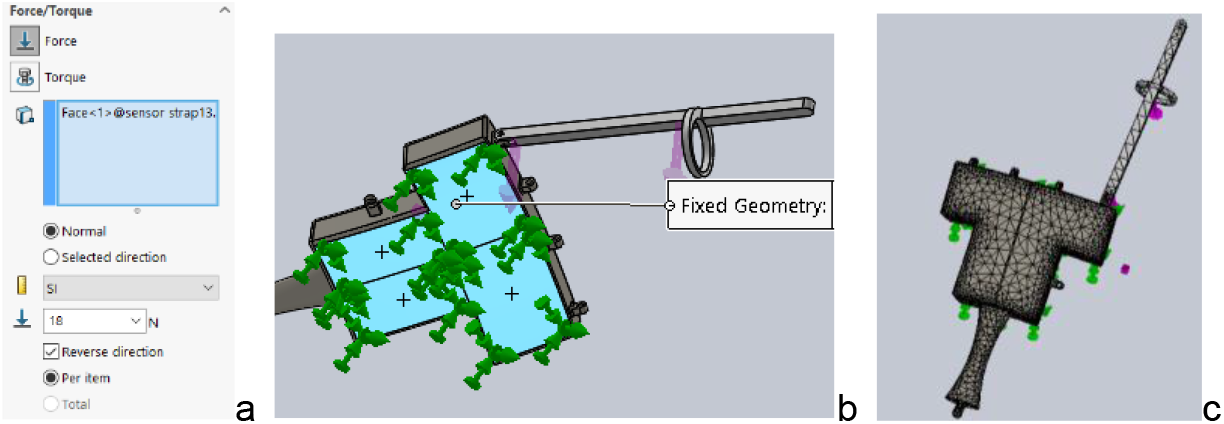
Introduced boundary conditions (a); fixture and Force setup (b); mesh overview (c)

**Figure 3.8:**
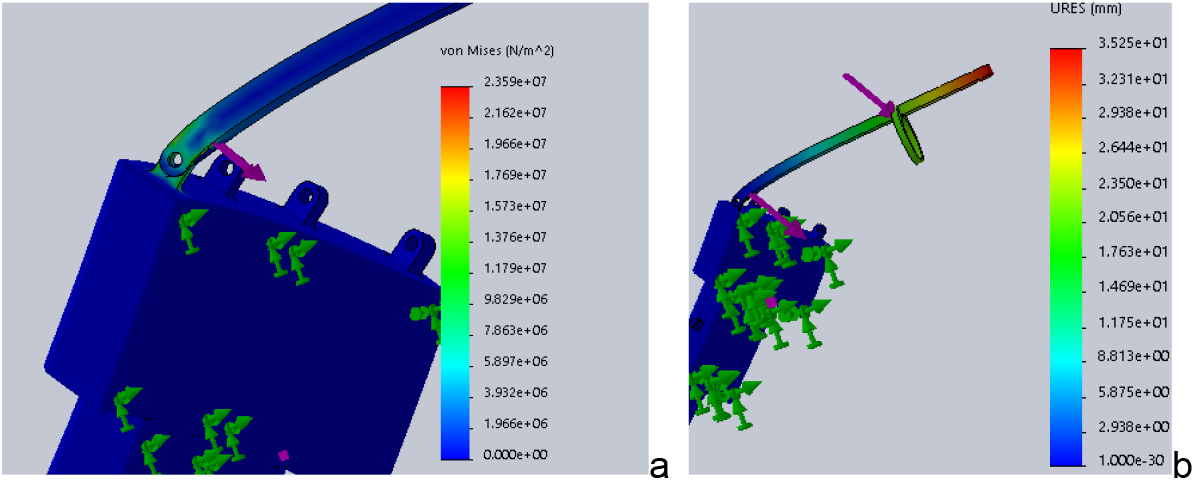
Simulation result: von Mises stress (a); displacement (b)

## 4 CONCLUSION AND RECOMMENDATIONS

In summary, the aim of designing a passive bionic grasper with basic performance and building up a virtual prototype of it has been accomplished in an iterative, time-efficient process. A thorough review of past literatures has provided sufficient information for generating the specific ideas and considerations for the design. A literature research has been accomplished for studying the background and developing the ideas of the design. Design concepts were processed based on PDS. In PDS process, 3 concept designs and components selection were presented and compared regarding to 6 parameters essential to a commercially viable smart grasper. The CAD model was built up and assembled to the hand model in SolidWorks. Drop simulation on the supporting pad and static evaluation applying loads on index finger strap were completed as well. The results showed that firstly, no failure would occur when the supporting pad was dropped from a 1.5m hight to the ground, no matter with the flat bottom face or the side edge of the smart grasper to touch the ground first, and secondly, yielding may appear on the structure employing concentric and coincident mates, which should be improved to avoid failure on the final product. Although, the results obtained here can be questioned due to the choice of simulation software, mesh quality, boundary conditions, time limitation, etc. The full validate of the results is possible only through a set of experimental examination. It is therefore suggested a few prototypes of the design be manufactured and a range of practical tests be applied on the prototyped samples.

## REFERENCES

1. Akpa, A., Fujiwara, M., Suwa, H., Arakawa, Y. and Yasumoto, K. (2019) A smart glove to track fitness exercises by reading hand palm, Journal of Sensors, 2019, 1–19.

2. Ben-Tzvi, P. and Ma, Z. (2015) Sensing and Force-Feedback Exoskeleton (SAFE) robotic glove, IEEE Transactions on Neural Systems and Rehabilitation Engineering, 23(6), 992–1002.

3. Bhardawaj, H., Dhaker, M., Sivani, K. and Vardhan, H. (2018) Smart Gloves: A novel 3-D Work Space Generation for Compound Two Hand Gestures, Proceedings of the 2018 International Conference on Control and Computer Vision, 28–32.

4. Böcker, S., and Malmström, S. (2018). Degree Project in Mechanical Engineering: Development of test system for soft robotic gloves.

5. Cerulo, I., Ficuciello, F., Lippiello, V. and Siciliano, B. (2017) Teleoperation of the SCHUNK S5FH under-actuated anthropomorphic hand using human hand motion tracking, Robotics and Autonomous Systems, 89, 75–84.

6. Chan, K., Ichikawa, K., Watashiba, Y. and Iida, H. (2017) Cloud-based VR gaming: our vision on improving the accessibility of VR gaming, International Symposium on Ubiquitous Virtual Reality, 24–25.

7. Cobos, S., Ferre, M., Uran, M., Ortego, J. and Pena, C. (2008) Efficient human hand kinematics for manipulation tasks, IEEE/RSJ International Conference on Intelligent Robots and Systems, 2246–2251.

8. Das, A., Yadav, L., Singhal, M., Sachan, R., Goyal, H., Taparia, K, Gulati, R., Singh, A. and Trivedi, G. (2016) Smart glove for sign language communications, 2016 International Conference on Accessibility to Digital World, 27–31.

9. Diftler, M., Bridgwater, L., Rogers, J., Laske, E., Ensley, K., Lee, J., Ihrke, C., Davis, D., and Linn, D.M. (2015) RoboGlove—A grasp assist device for earth and space, In Proceedings of the 45th International Conference on Environmental Systems, 1–6.

10. Dipietro, L., Sabatini, A. and Dario, P. (2008) A survey of glove-based systems and their applications, IEEE Transactions on Systems Man and Cybernetics Part C: Applications and Reviews, 38, 461–482.

11. Francés, L., Morer, P., Rodriguez, M. I., & Cazón, A. (2019). Design and Development of a Low-Cost Wearable Glove to Track Forces Exerted by Workers in Car Assembly Lines. Sensors (Basel, Switzerland), 19(2), 296.

12. Frawan, L., Basmaji, T. and Hassanin, O. (2018) A mobile mental health monitoring system: a smart glove, 2018 14th International Conference on Signal-Image Technology & Internet-Based System, 235–240.

13. Gu, X., Zhang, Y., Sun, W., Bian, Y., Zhou, D. and Kristensson, P. (2016) Dexmo: An inexpensive and lightweight mechanical exoskeleton for motion capture and force feedback in VR, Proceedings of the 2016 CHI Conference on Human Factors in Computing Systems, 1991–1995.

14. Heo, P., Gu, G., Lee, S., Rhee, K. and Kim, J. (2012) Current hand exoskeleton technologies for rehabilitation and assistive engineering, International Journal of Precision Engineering and Manufacturing, 13(5), 807–824.

15. Hirata, Y., Tozaki, Y. and Kosuge, K. (2011) Development of wire-type motion support system controlled by Servo Brakes, Proceedings of the 2011 IEEE International Conference on Robotics and Biomimetics, 2150–2155.

16. Hosseini, M., Sengul, A., Pane, Y., Schutter, J. and Bruyninckx, H. (2018) ExoTen-Glove: A force-feedback haptic glove based on twisted string actuation system, Proceedings of the 27th IEEE International Symposium on Robot and Human Interactive Communication, 320–327

17. In, H., Cho, K., Kim, K., and Lee, B. (2011) Jointless structure and under-actuation mechanism for compact hand exoskeleton. In: 2011 IEEE international conference on rehabilitation robotics (ICORR), Zurich, 29 June–1 July, 1–6.

18. Jeong, U. and Cho, K. (2016) A novel low-cost, large curvature bend sensor based on a Bowden-cable, Sensors, 16(961), 1–20.

19. Jung, H., Kim, H., Jeong, J., Jeon, B., Ryu, T. and Kim, Y. (2017) Feasibility of using the RAPAEL Smart Glove in upper limb physical therapy for patients after stroke: A randomized controlled trial, Conference proceedings: Annual International Conference of the IEEE Engineering in Medicine and Biology Society, IEEE Engineering in Medicine and Biology Society. Annual Conference, 2017, 3856–3859.

20. Kortier, H. Sluiter, V., Roetenberg, D. and Veltink, P. (2014) Assessment of hand kinematics using inertial and magnetic sensors, Journal of NeuroEngineering and Rehabilitation, 11(70), 1–14.

21. Kortier, H., Schepers, H. and Veltink, P (2016) Identification of object dynamics using hand worn motion and force sensors, Sensors, 16, 1–17.

22. Lariviere, C., Tremblay, G., Nadeau, S., Harrabi, L., Dolez, P., Vu-Khanh, T. and Lara, J. (2010) Do mechanical tests of glove stiffness provide relevant information relative to their effects on the musculoskeletal system? A comparison with surface electromyography and psychophysical methods, Applied Ergonomics, 41, 326–334.

23. Li, X., Wen, R., Shen, Z., Wang, Z., Luk, K. and Hu, Y. (2018) A wearable detector for simultaneous finger joint motion measurement, IEEE Transactions on Biomedical Circuits and Systems, 12(3), 644–654.

24. Lin, B., Lee, I., Chiang, P., Huang, S. and Peng, C. (2019) A modular Data Glove system for finger and hand motion capture based on inertial sensors, Journal of Medical and Biological Engineering, 39, 532–540.

25. Ma, Z. and Ben-Tzvi, P. (2015) RML Glove—An exoskeleton glove mechanism with haptics feedback, IEEE/ASME Transactions on Mechatronics, 20(2), 641–652.

26. Placidi, G. (2007) Asmart virtual glove for the hand telerehabilitation, Computers in Biology and Medicine, 37, 1100–1107.

27. Salchow-hommen, C., Callies, L., Laidig, D., Valtin, M., Schauer, T. and Seel, T. (2019) A tangible solution for hand motion tracking in clinical applications, Sensor, 19(208), 1–27.

28. Shahid,T., Gouwanda, D., Nurzaman, S. and Gopalai, A. (2018) Moving toward soft robotics: A decade review of the design of hand exoskeletons, Biomimetics, 3(17), 1–20.

29. Silva, L., Dantas, R., Pantoja A. and Pereira, A. (2013) Development of a low cost dataglove based on arduino for virtual reality applications, 2013 IEEE International Conference on Computational Intelligence and Virtual Environments for Measurement Systems and Applications (CIVEMSA), Milan, 2013, 55–59.

30. Shin, J., Kim, M., Lee, J., Jeon, Y., Kim, S., Lee, S., Seo, B., and Choi, Y. (2016). Effects of virtual reality-based rehabilitation on distal upper extremity function and health-related quality of life: a single-blinded, randomized controlled trial, Journal of neuroengineering and rehabilitation, 13 (17), 1–10.

31. Wang, D., Guo, Y., Liu, S., Zhang, Y., Xu, W. and Xiao, J. (2019) Haptic display for virtual reality: progress and challenges, Virtual Reality & Intelligent Hardware, 1(2), 136–162.

32. Weber, P., Rueckert, E., Calandra, R., Peters, J. and Beckerle, P. (2016) A low-cost sensor glove with vibrotactile feedback and multiple finger joint and hand motion sensing for human-robot interaction, 2016 25th IEEE International Symposium on Robot and Human Interactive Communication (RO-MAN), 99–104.

33. Wilk, M., Torres-Sanchez, J., Tedesco, S. and O’Flynn, B. (2018) Wearable human computer interface for control within immersive VAMR gaming environments using Data Glove and hand gestures, IEEE Games, Entertainment, Media Conference (GEM), 213–219.

34. Yu, A., Yick, K., Ng, S., and Yip, J. (2013). 2D and 3D anatomical analyses of hand dimensions for custom-made gloves. Applied ergonomics, 44(3), 381–392.

35. Zheng, J., Rosa, S. and Dollar, A. (2011) An investigation of grasp type and frequency in daily household and machine shop tasks, IEEE International Conference on Robotics and Automation, 4169–4175.

36. Zheng,Y., Wang, D., Wang, Z., Zhang, Y., Zhang, Y. and Xu, W. (2018) Design of a lightweight force-feedback glove with a large workspace, Engineering, 4, 869–880.

37. Zou, Y., Wang, D., Hong, S., Ruby, R., Zhang, D. and Wu, K. (2020) A low-cost smart glove system for real-time fitness coaching, IEEE Internet of Things Journal, 1–16

